# Toblerone: detecting exon deletion events in cancer using RNA-seq

**DOI:** 10.1101/2022.10.27.514132

**Authors:** Andrew Lonsdale, Andreas Halman, Lauren M Brown, Hansen J Kosasih, Paul G Ekert, Alicia Oshlack

## Abstract

Cancer is driven by mutations of the genome that can result in the activation of oncogenes or repression of tumour suppressor genes. In acute lymphoblastic leukemia (ALL) focal deletions in IKAROS family zinc finger 1 (IKZF1) result in the loss of zinc-finger DNA-binding domains and a dominant negative isoform that is associated with higher rates of relapse and poorer patient outcomes. Clinically, the presence of IKZF1 deletions informs prognosis and treatment options. In this work we developed a method for detecting exon deletions in genes using RNA-seq with application to IKZF1. We developed a pipeline that first uses a custom transcriptome reference consisting of transcripts with exon deletions. Next, RNA-seq reads are mapped using a pseudoalignment algorithm to identify reads that uniquely support deletions. These are then evaluated for evidence of the deletion with respect to gene expression and other samples. We applied the algorithm, named Toblerone, to a cohort of 99 B-ALL paediatric samples including validated IKZF1 deletions. Furthermore, we developed a graphical desktop app for non-bioinformatics users that can quickly and easily identify and report deletions in IKZF1 from RNA-seq data with informative graphical outputs.

## Introduction

B-cell precursor ALL (BCP-ALL, or B-ALL) is the most common childhood cancer. High risk subtypes of B-ALL include Ph+ (presence of BCR-ABL1 fusion) and Ph-like (similar expression profile to Ph+ without BCR-ABL1 fusion) (Roberts et al. 2014). These subtypes are often characterised by additional focal deletions in IKAROS family zinc finger 1 (IKZF1) gene. IKZF1 consists of 8 exons and encodes a DNA-binding protein and B-cell transcription factor, IKAROS (Mullighan et al. 2009; Boer et al. 2016). The most common IKZF1 alterations that occur in B-ALL are whole gene deletions or deletions of exons 4-7. The focal deletion results in the production of a dominant-negative isoform, IK6, lacking the N-terminal DNA-binding domains of IKAROS (Mullighan et al. 2009; Dörge et al. 2013; van der Veer et al. 2013). Previously we and others have showed that IK6 could be detected using RNA-seq (Brown et al. 2020; Tran et al. 2022; Rehn et al. 2022). In our work, we did so by adding the deletion transcript to the reference transcriptome and measuring the transcripts per million (TPM) (Brown et al. 2020). Given IK6 lacks exons 4-7 (del4_7) of the canonical IKZF1 transcript, this isoform includes a novel splice junction between exon 3 and exon 8. In our cohort we found that the splice junction indicating an exon deletion was rarely detected in the cohort in IK6-negative samples, and was increasingly expressed as the purity of the tumour increased as verified by real-time quantitative polymerase chain reaction (RQ-PCR) (Brown et al. 2020). Other isoforms of IKZF1 deletions in the cohort were not as reliably detected. Only one sample was known to have a deletion of exons 2-7 (del2_7) and our previous predictions of increased expression of the del2_7 transcript were unable to be validated. Previous candidates of deletions of exons 2-8 (del2_8) and exons 4-8 (del4_8) were also unable to be reliably validated. Here we develop an improved method for detecting focal deletions that can be applied more broadly to other gene deletions. Prior knowledge of which exon deletions are of interest is not required.

The key idea in our method is to generate a set of reference transcripts that correspond to all the possible exon deletions in a gene. This idea represents something of a midpoint between using annotated transcripts as the reference transcriptome and using assembly, by pre-defining potential transcripts in the class of whole exon skipping. Essentially, we generate a reference index of possible transcripts based solely on exon deletions of annotated exons, balancing the speed of a reference based approach while allowing for a certain class of unknown events. We extract reads that support these deletions, test for sufficient coverage (expression) of a deletion, and then infer the relative abundance of deletion reads within a gene.

We name this method Toblerone and apply it to a B-ALL cohort from the Royal Children’s Hospital (RCH), previously described in Brown et al. We validate the method on the IKZF1 gene and identify additional deletions with high relative expression of the deletion transcript. Toblerone can be applied in both a test and discovery context. Firstly, it can be used to test for known deletion events such as IK6 (exons 4-7 deletion) in IKZF1 or secondly, it can be used for discovery of new genes with exon deleted transcripts, by applying it to any gene with three or more exons. Toblerone is a tool that can quickly and accurately identify skipped exons that may indicate the presence of sub-gene deletions in a cancer sample.

## Results

### Toblerone method overview

Toblerone begins by creating a custom reference. For any candidate gene, e.g. IKZF1, we take the canonical transcript and generate a new transcriptome reference that consists of the original transcript plus a set of deletion transcripts. Deletion transcripts consist of removing combinations of consecutive internal exons, excluding edge exons (first and last). This avoids the lack of splicing events at the ends of genes. A schematic of deletion transcripts for a five exon gene is shown in Figure 1A, resulting in six deletion transcripts.These deletion transcripts along with the canonical transcript are used as the complete reference for the gene. The new custom reference is then indexed using a De Bruijn graph for pseudoalignment of RNA-seq reads.

**Figure 1.**
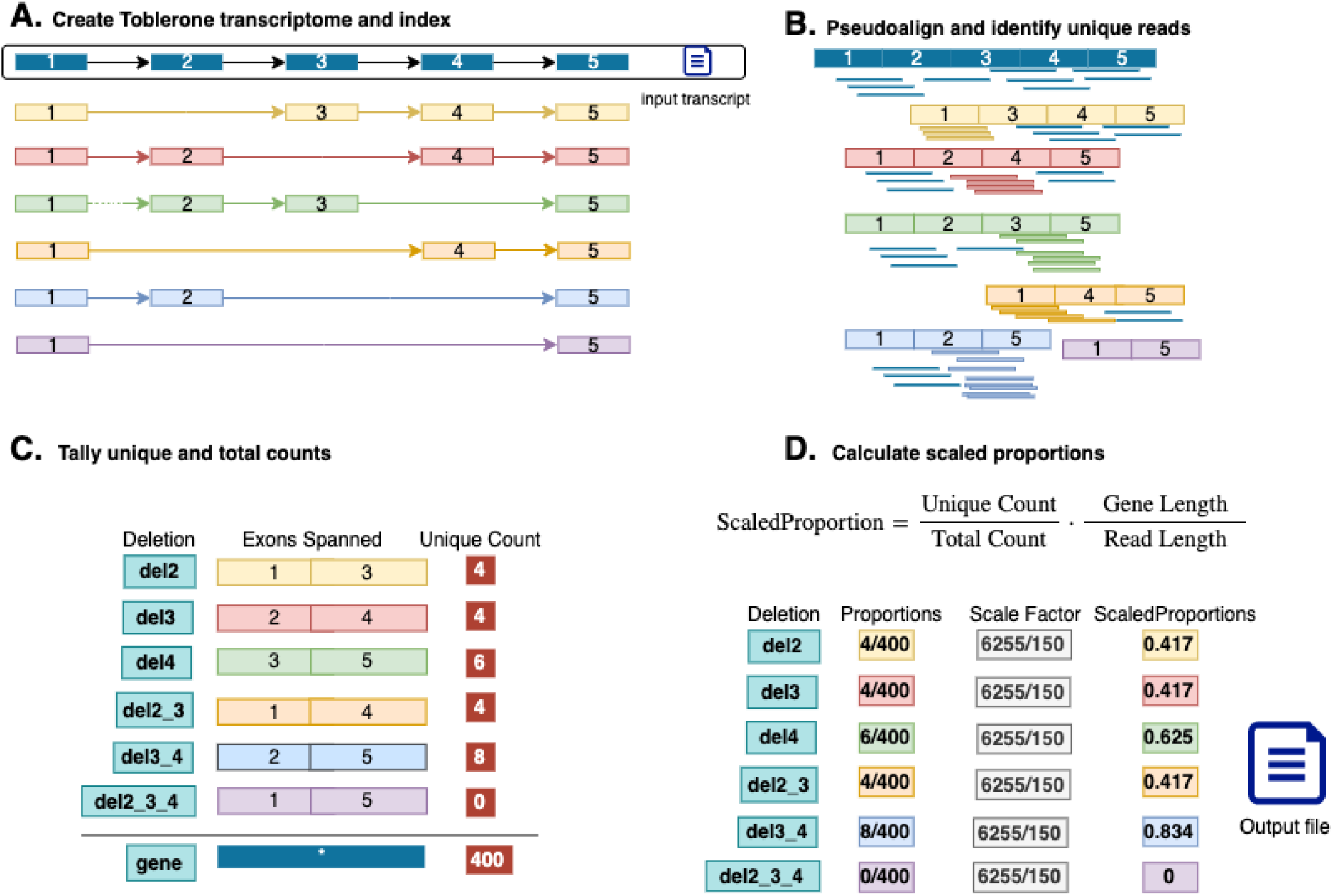
Overview of the Toblerone method. A) Using a canonical input transcript, deletion transcripts are generated and indexed into a custom reference. B) Reads are then pseudoaligned to the custom reference. C) reads uniquely supporting a deletion transcript are aggregated into the unique count (UC) for each transcript. All other reads are added to gene total. D) Scaled proportions of unique counts (UC) to totals counts (TC) are calculated for each deletion and saved to the output file.

Next reads are pseudoaligned to the custom reference and quantified at the equivalence class (EC) level based on which of the transcripts they are compatible with. Due to the mutually exclusive construction of the deletion transcripts, there is one EC that is uniquely compatible with each deletion. These correspond to the reads that have splice junctions across the deleted exons. Rather than using the transcript-compatibility count (TCC) (Ntranos et al. 2016) of a deletion transcript, derived from all EC that would support a deletion transcript, the Toblerone approach uses only the count from the EC uniquely supporting that deletion (Figure 1B). We refer to the EC counts for this subset of EC as the unique count (UC) of each deletion transcript. The reads for the remainder of EC, those that support the original canonical transcript for a gene, or support more than one deletion transcript, are used in the sum total of EC counts as an expression measure for the gene. We refer to this as the total EC count for a gene, or simply the total count (TC). Deletion detection requires coverage across the junction including overhang of the exon-exon boundary by at least 5 bp (by default). Any read that does not meet this threshold is not included as a UC, only in the TC. Deletion UC as a proportion of the TC is calculated and scaled by the length of the canonical transcript and the read length to account for the increased number of reads overlapping a splice junction with increased read length. A command line tool is available to generate and index the deletion transcriptome, and perform pseudoalignment. See Methods for further details.

### Application to paediatric acute lymphoblastic leukemia cohort

We applied Toblerone to the IKZF1 gene from RNA-seq data from a cohort of 99 B-ALL patients (Figure 2), which included annotations for validated IKZF1 deletion status from microarray and RQ-PCR, as well as other clinical data including fusion status and karyotyping result (Brown et al. 2020). The scaled proportions for each of the 21 IKZF1 deletion transcripts are shown in Figure 2A, and illustrate the different frequencies of exon deletions. The majority of deletions are rarely observed in the cohort, with several notable exceptions. Deletions of exon 4 and exon 6 are commonly observed and are consistent with IKZF1 annotated alternative splicing transcripts attributed to hg38 reference transcripts (ENST00000343574, ENST00000359197, ENST00000439701). The relative expression of these transcripts is variable across samples but does not usually account for more than half of the transcripts based on the calculated proportions. For IK6, the validated del4_7 deletions have proportions that make them clearly identifiable visually, a result consistent with the previous methods applied to this cohort. In order to test for outliers in the proportion of reads indicating a deletion, we transformed the scaled proportion into a Z-score for each deletion. The distribution of z-scores for deletion transcript with at least one sample with Z-score greater than or equal to three is shown, along with its outliers (Figure 2B). Since low gene expression can lead to high Z-score and therefore outliers with only a small number of counts, we exclude deletions that had outliers due to samples with UC counts of 10 or less. These are however included in Supplementary Figure 1. The validated del4-8 sample is also observed as an outlier of exon 4 deletions. In this case, the signal is indirect as loss of half the exons alters the proportion of splicing events of exon4 relative to the gene expression. Outlier detection identifies two other known IKZF1 deletions; the single known del2_7 deletion in the cohort and a new result of a del3_7 deletion previously only predicted from a low resolution microarray.

**Figure 2.**
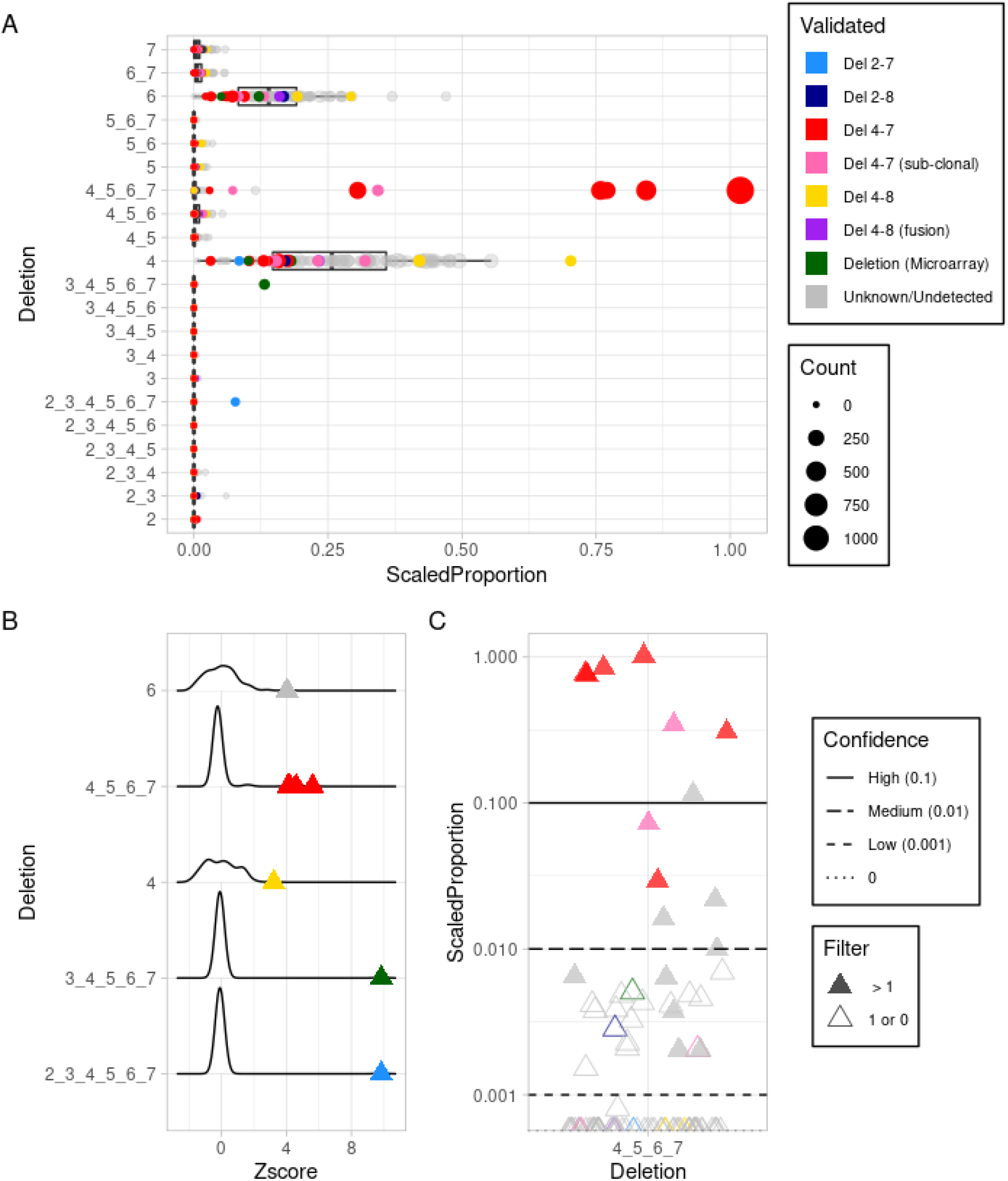
Application of Toblerone to RCH cohort. A) Boxplots of scaled proportions of unique counts (UC) for each transcript of IKZF1 exon deletions. Each row is labelled with the exon numbers that are deleted in that transcript. Each sample is a dot with a size proportional to the UC and coloured by known IKZF1 deletion status (light blue: del2_7, dark blue: del2_8, red: del4_7, pink: del4_7 sub-clonal, yellow: del4_8, purple: del4_8 fusion, green: non specific deletion from DNA microarray, grey: deletion unknown or undetected). B) Distributions of the scaled proportions transformed into a Z-score within each deletion. Only the IKZF1 deletion transcripts with at least one Z-score outlier ≥ 3 and minimum UC count of 10 are shown, with outliers shown as triangles coloured as in A. C) Scaled proportion scores of IKZF1 del4_7 for the cohort with high, medium and low confidence lines at 0.1 (solid), 0.01 (longdash) and 0.001 (dashed). Samples with two or more counts of the IK6 UC are filled triangles while zero or one count is empty. Many samples including sub-clonal true positives have 0 counts (dotted line).

Although outliers can easily identify four of the six IK6 deletions, identifying the remaining two clonal, as well as a further five sub-clonal deletions presents additional challenges. In Figure 2C we show the calculated del4_7 proportions for every sample in the cohort on a log scale. We again observe the high proportion of IK6 UC as outliers and can empirically establish thresholds for high, medium and low confidence calls for the IKZF1 del4_7 deletion. We define high confidence at 0.1 scaled proportion, as most values exceeding this are known IK6 deletions; medium is set at a scaled proportion of 0.01 and a low confidence threshold of 0.001 isolates a cluster of samples that include some sub-clonal IK6 or unresolved microarray deletions as well as many unvalidated samples. The choices of these thresholds are informed by a ROC curve of validated true positives and negatives (Supplementary Figure 2). The area under the ROC of 0.859 confirms that Scaled Proportions effectively predict true positive del 4-7 samples, and the thresholds to identify when false-positives may be introduced. In the low confidence band especially, many samples have a single read supporting the del4_7 deletion. Low or singular counts are insufficient to identify deletions, and two validated sub-clonal IK6 samples lacked any reads supporting deletion and had counts of 0. Inclusion of the low confidence results may lead to increased false positives due to the possibility of technological or biological artefact, and so would not typically be recommended.

### Effect of parameters on results

EC counts for Toblerone deletions can be heavily influenced by the alignment of reads across a junction which is controlled by parameters set at runtime, specifically mismatch and trim. The mismatch parameter controls how many single nucleotides each kmer can differ from the reference in order to match. Since reads at exon-exon boundaries form the signal of Toblerone, reads that overhang one exon by a small number of bases, equal to the number of allowed mismatches, may be erroneously assigned as supporting a deletion. Conversely disallowing mismatches can reduce the signal from genuine deletion by excluding any reads that contain genomic variants or mutations. To balance the effects of allowing mismatches, Toblerone includes a trim operator which tests the robustness of read support. For each read that uniquely supports a deletion, nucleotides are removed from the ends of the read and the support checked again. Reads that are still unique and only support the deletion pass the trim test. Otherwise these reads contribute only to the gene total. The trim operation essentially prevents soft-clipped reads contributing to the UC total, and reduces noise by removing reads with a small overhang of bases at the exon-exon boundary, ensuring that any mismatches are not the sole evidence a read supports a deletion. The trim value must equal or exceed the mismatch value. The effect of Toblerone’s trim and mismatch on the counts, and the effect of the changes in these counts to the scaled proportion, is illustrated in Supplementary Figure 3A and 3B respectively.

The total number of samples with any read support for IKZF1 del4_7 is shown in Supplementary Figure 3A for combinations of trim and mismatch. With default parameters, 34 samples could be detected with one or more reads. If both trim and mismatch are set to 0, this reduces to 31 samples. The number of samples with observed del4_7 deletion reads are typically reduced by higher trimming values, and increased by allowing more mismatched bases. Although the change in the number of samples is modest, the effect of these parameters is observable in the calculated scaled proportions in some samples (Supplementary Figure 3B). Though the majority of samples are relatively unchanged, notably two validated deletions have an increased proportional expression when including trimming and mismatches.

### Toblerone app for single sample analysis

In addition to a command line tool, a Toblerone graphical desktop app for non-bioinformatics users was created (Figure 3A). This enhances the command line tool by allowing a user to provide either a hg19 or hg38 genome aligned BAM file. Relevant reads for IKZF1 are extracted and then pseudoaligned against the Toblerone IKZF1 transcriptome using the core Toblerone command line tool that has been integrated into the app. The results are then displayed and compared against the background cohort. The app uses the RCH cohort explored above, as a background, and the same parameters are used as defaults when comparing to new data. The high, medium and low thresholds established for this cohort are also included in the app. We demonstrate the app using a relapse sample from a patient in the RCH cohort (B_ALL16-4). The primary sample was validated as a sub-clonal IKZF1 del4_7 deletion and included in the cohort. The relapse sample was subsequently validated as a full IKZF1 deletion, though not included in the cohort data in Toblerone. The result of submitting this RNA-seq sample within the Toblerone app is shown in Figure 3. Firstly, the app indicates that the del4_7 deletion has been detected in the sample, and assigns it within the high confidence band (Figure 3A) with a Scaled Proportion of 0.949. Z-scores for this sample for all deletions in IKZF1 are shown in Figure 3B, and outliers are indicated in red. The app also gives details for each deletion in a separate tab, with the background deletion distribution able to be inspected. This is shown for IK6 and the relapse sample in Figure 3C, where it is clear the Scaled Proportion of the relapse sample is greater than any value seen in the background cohort. This plot can also be viewed as Z-scores if selected by the user. See Methods for further installation and usage information for the Toblerone application.

**Figure 3.**
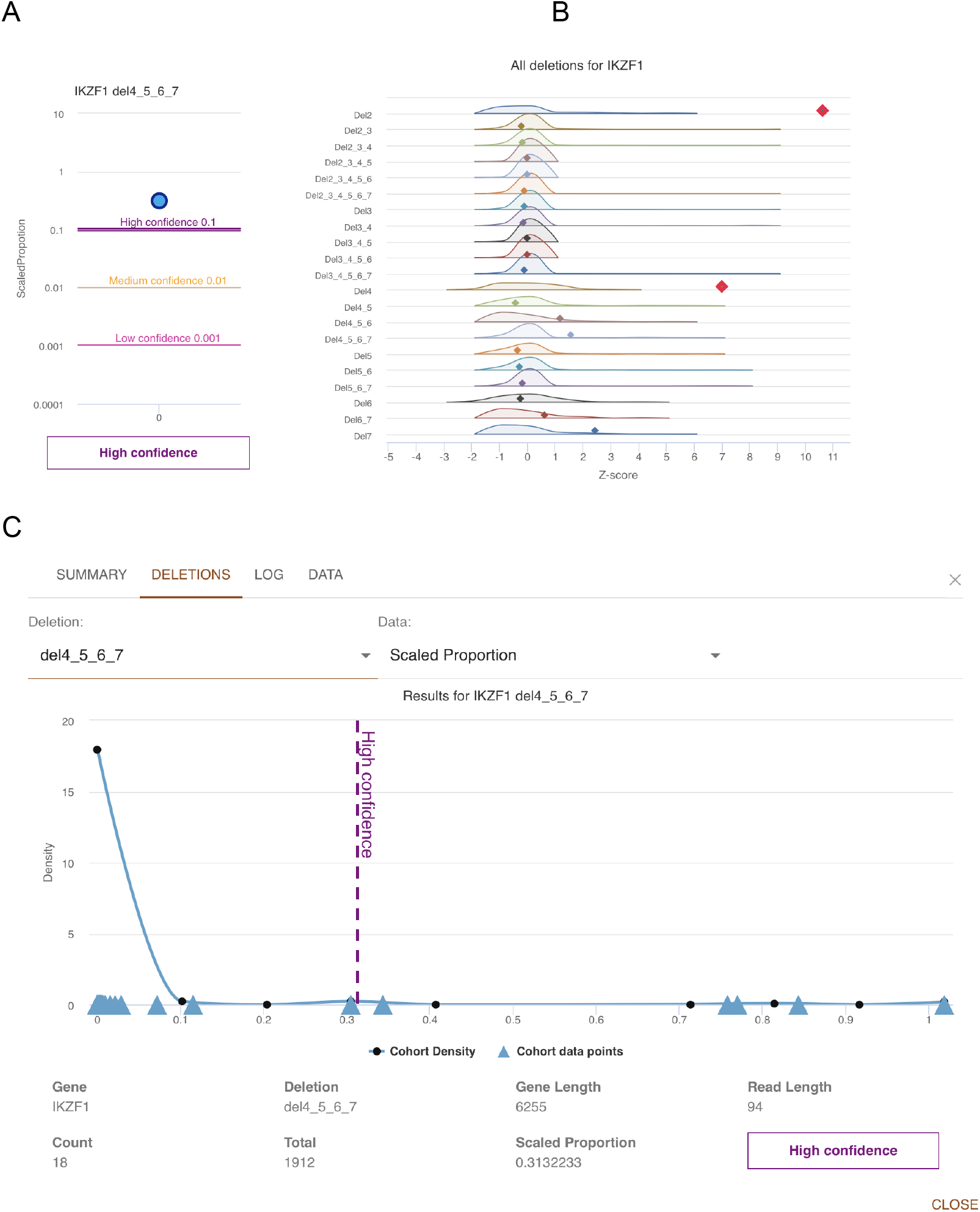
Overview of Toblerone app applied to a new sample. A) Scaled proportion of the IKZF1 del4-7 compared to the previously calculated confidence thresholds; B) the Z-score distributions, with the sample indicated by dots for all IKZF1 deletion transcripts. Any outliers are shown in red; C) detailed view of the del4-7 background distribution in the RCH cohort, with gaussian density estimate line in blue and each cohort value as triangles along the x-axis. The B_ALL16_4 sample is indicated by vertical dotted line, coloured by the confidence value.

### Application to CD22 resistance in CAR T-cell therapy

While Toblerone has been developed and tested on the IKZF1 genes across a cohort of samples it can also be used with different genes and to compare between samples in paired study design. As a proof of concept, we use data from a recent publication exploring acquired resistance when using CD22 as an immunotherapeutic target for CAR T-cell therapy (Zheng et al. 2022). In that work, RNA-seq data was taken before and after for B-ALL patients undergoing inotuzumab treatment. Notably for the patient with specimen identifier PAVDRV, a decrease in CD22 protein expression after treatment occurred without down regulation of the CD22 transcript. To compare Zheng’s findings with a Toblerone approach, a CD22 index was created and Toblerone was applied to the samples pro- and post-treatment (no replicates). We were able to confirm the study’s result that a transcript with loss of exon 2, specifically the deletion of exons 2-6, was increased after treatment with inotuzumab (Figure 4). Additionally, we ranked and visualised the Toblerone differences in deletion proportions of all transcripts in the Toblerone reference (Supplementary Figure 3). This showed that post-inotuzumab treatment there is an increase in expression of the transcript with an exon 12 deletion. While the original publication noted this as a known deletion observed in the B-ALL samples of the TARGET cohort, it was not attributed to patient PAVDRV as a possible cause of the loss of CD22 protein.

**Figure 4.**
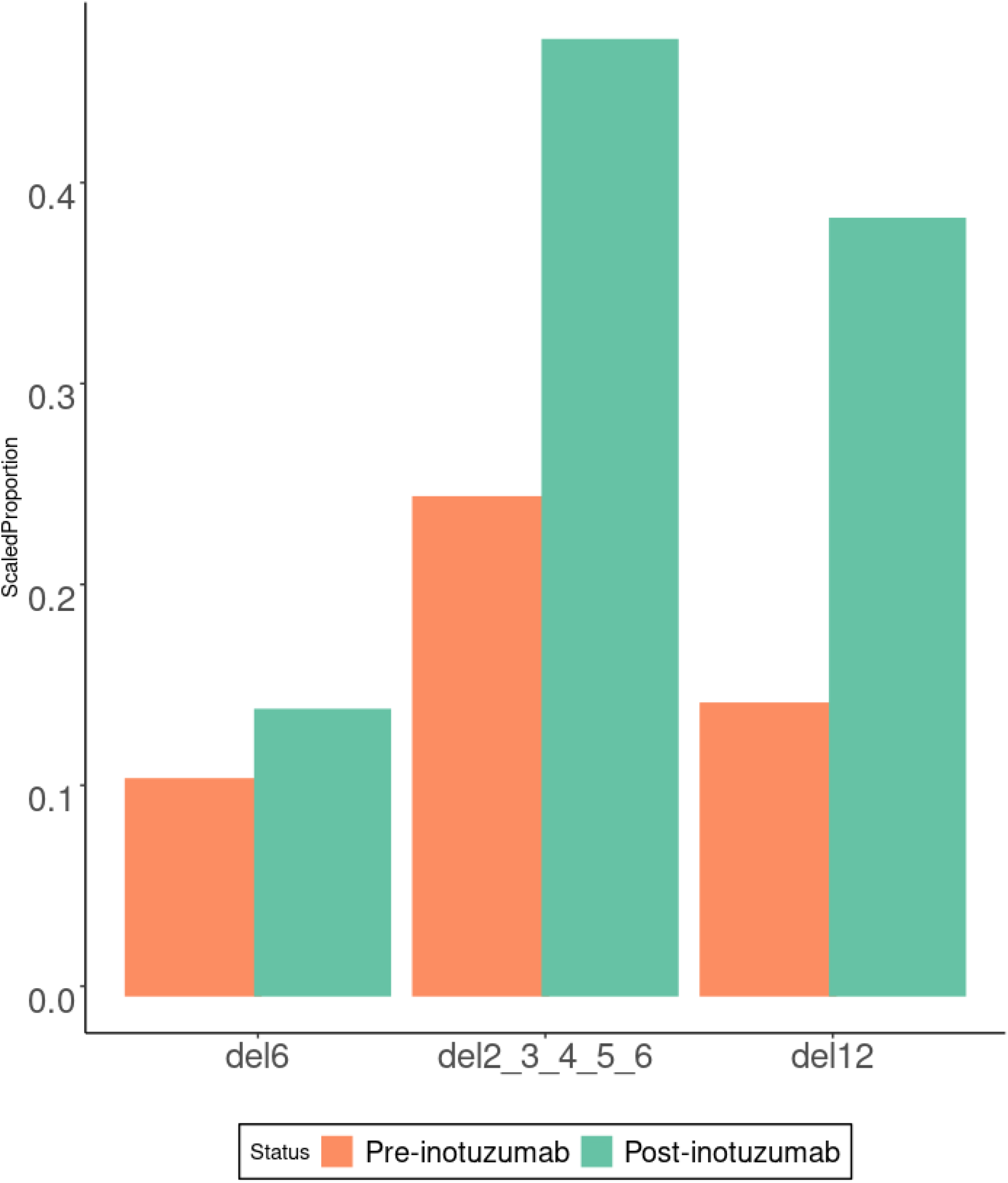
Differences in scaled proportions of the 3 most highly expressed CD22 deletion transcripts for patient identifier PAVDRV. Coloured bars represent the patient sample taken from peripheral blood before (orange) and after (green) treatment with inotuzumab. Selected deletions showed most deviation from pre-treatment results (Supplementary Figure 3).

## Discussion

Toblerone is a targeted technique for identifying internal exon deletions in specific genes from RNA-seq. Inspired by the detection of deletions in the IKZF1 gene in Brown et al, we have tailored an algorithm to specifically look for these novel isoforms in RNA-seq. A recent approach that performs more comprehensive RNA-seq analysis specific to ALL is RasCALL (Rehn et al. 2022). Toblerone is not intended to be a comprehensive structural variant detection method however it complements existing RNA-seq analysis tools in cancer such as fusion detection (e.g. JAFFA (Davidson, Majewski, and Oshlack 2015), Aribba (Uhrig et al. 2021), STAR-fusion etc) and SV detection (e.g. MINTIE (Cmero et al. 2020)). These and other tools may be more suitable for broader detection of unusual fusions, transcripts or more complex deletions, especially in situations where atypical events may be missed by standard of care molecular diagnostics (Nardi et al. 2022). Similarity, Toblerone is not intended for comprehensive analysis of alternative splicing from RNA-seq. LeafCutter (Y. I. Li et al. 2018) and MAJIQ (Vaquero-Garcia et al. 2016) may be more suitable when considering these changes at a whole transcriptome level.

The Toblerone analysis has been clearly demonstrated for the detection of deletions in IKZF1 including the most commonly seen deletion involving exons 4 to 7 as well as rarer deletions with other combinations of exons. The algorithm can also be extended to discover novel exon deletions in other genes, with some caveats. Toblerone is only applicable to genes with three or more exons where the deletion does not involve the loss of the terminal exons. Loss of terminal exons could potentially be seen in differential expression analysis at the exon or gene level. If these conditions are met, then parallel indexes for additional genes of interest for a given cohort could be executed, e.g. PAX5 in B-ALL (Mullighan et al. 2009). Toblerone indexes can also be created for other organisms, following the same index creation process with substituted reference files and gene definitions. In IKZF1 research mouse models of Ikzf1 can be used to understand its role in immunity and diseases related to Ikaros (Boast et al. 2021). Example indexes for PAX5 (hg38) and Ikzf1 (mm10) are available with the Toblerone source code.

When a deletion transcript is lowly expressed it is difficult to detect with RNA-seq. Toblerone uses junction read counts to infer the novel junctions produced by a deletion. Even with checks to ensure that reads supporting deletions are not due to minor overlaps at exon boundaries, singular reads for deletions were found in many samples without validated deletions. Conversely, we were not able to detect any supporting reads in several subclonal del4_7 deletions. Determining whether these low counts are a biological or technical artefact remains an open question. With this in mind we proposed thresholds for high, low and medium confidence deletions to provide a guide to interpreting Toblerone results, as well as establishing the level of background noise for the method. Extending to other genes and cohorts may require validated samples to establish appropriate thresholds, between validated samples and remainder of the cohort samples. However, as shown with acquired resistance with CD22 case study, albeit without sufficient replicates for statistical analysis, Toblerone can also be used in case-control studies to compare exon deletions proportions between two groups.

The generation of the Toblerone transcriptome and pseudoalignment is targeted towards single gene analysis and therefore is not optimised for a complete transcriptome or competitive pseudoalignment between multiple genes. Results presented here have used extracted reads from genome alignments, and as such the reads have already been aligned by more conventional approaches such as STAR (Dobin et al. 2013). Conceptually it is possible to use raw reads and implement the Toblerone approach as part of a competitive pseudoalignment, for single or multiple genes of interest. The Toblerone custom deletion transcriptome can be used with any pseudoaligner that divulges EC counts prior to abundance estimation. So conceptually a complete pseudoalignment from raw FASTQ data can be performed in programs such as Kallisto (Bray et al. 2016) and Salmon (Patro et al. 2017) with a mixture of reference transcripts and additional deletions transcripts. This approach however stores EC counts compatible with more than one transcript, which is superfluous for the needs of the Toblerone algorithm, and this additional overhead increases with each gene considered and prevents efficient computation across multiple samples. The Toblerone pseudoaligner is designed to only compute the necessary counts, and is optimised to identify reads that support deletions. Future work on Toblerone however can address these limitations for multi-gene analysis in a competitive alignment, through improvements to the Toblerone pseudoaligner or by reprocessing conventional pseudoaligner results to reduce redundant counts, and removing reads that do not robustly support a deletion.

The fundamental idea of Toblerone is to modify the transcriptome reference to contain transcripts with all variations for a sequence event of interest. By generating a transcriptome that consists of transcripts with deleted exons, reads that uniquely support that transcript can be identified, inspected and compared. Here, we have modified the reference to represent exon deletions, but there are other possible variants which may be detected. For example, intron retention is a natural extension of the existing algorithm, and exon duplications or inversions could also be identified. Toblerone adds insight into the clinically relevant transcriptional consequences of mutations by extending the analysis of RNA-seq, which is being applied more generally in many malignancies, notably lymphoblastic leukaemia. It can quantify the relative expression of known clinically relevant exon deletions in cancer, and aid in the discovery of new ones.

## Methods

### Toblerone index

The Toblerone transcriptome is generated in several steps, starting from a BED12 definition of the canonical gene(s) of interest that is given to the provided “make_bedfiles.py” Python script. New BED definitions for each Toblerone deletion transcript are then created. For each BED file entry of a transcript of length N exons, the number of deletions transcripts added for a given transcript can be determined using binomial theorem. For N exons, initially N-2 is used to calculate the internal combinations of exon deletions. The method requires continued runs of exons, coincidentally forming a pattern of triangular numbers. These can also be calculated binomially and equal to the binomial coefficient of t+1 and 2 for each t triangle number. Substituting N-2 for t leads to a binomial coefficient of *N*-1 and 2 for calculating the added Toblerone deletion transcripts. The results of this script are multiple BED files, which are merged and along with a FASTA file of the canonical transcripts corresponding to the BED12 input, passed to the bedtools (Quinlan 2014) program to create the Toblerone transcriptome FASTA file for indexing.

The core Toblerone program is an adaptation of a pseudoaligner written in the Rust programming language. This simple pseudoaligner, modified from 10XGenomics, implements concepts from several notable pseudoalignment and RNA-seq publications (Bray et al. 2016; Srivastava et al. 2016; Ntranos et al. 2016; Srivastava et al. 2019; Limasset et al. 2017; Orenstein et al. 2017; Y. Li and XifengYan 2015). Two main modes of operation are available: index and map. Index is used to create the de Bruijn index of the Toblerone transcriptome, provided a FASTA file of a generated transcriptome and an output file to store the index.

### Toblerone mapping

Map mode is used with either single or paired end reads as input, along with an index, to produce output. For optimal performance, only reads that are likely to match to genes of interest in the input transcriptome should be provided, such as by extracting reads from a prepared genome alignment. Toblerone diverges from the traditional pseudoalignment and template code in several critical ways during the mapping stage. Firstly, for any read the numbers of transcripts in the equivalence class is checked; any read matching an EC unique to one transcript is treated differently due to the mutually exclusive nature of the input transcriptome. The coverage of these reads over the input transcript must be complete, expecting the number of allowed mismatches (default: 2). These reads with a unique EC (if trim is enabled, default: 5) are checked for equality between the EC and the EC of a shorter trimmed version of the read.

Secondly, uniqueness is favoured when resolving ambiguity. When comparing the equivalence classes of paired-end reads, the matching EC with fewer transcripts is preferred in such cases ensuring that the read supporting a deletion is used in further processing. Thirdly, only unique EC (UC) are tallied individually in memory against a specific transcript and written to output, with all other reads compatible with more than one transcript assigned only to a gene-level total count (TC). For each deletion the UC and TC are written to file along with the gene length, nominated read length. The proportion of UC to TC for each deletion-gene combination is also included, as a raw value and with a scaling factor equal to the the gene length divided by the read length (paired end reads weighted double). The scaled proportions are then calculated by multiplying the scale factor by the original proportion (Figure 1). These results are written to a CSV file, or standard output, as directed by the user.

### Toblerone app

A graphical Toblerone app is written in Python and JavaScript programming languages, and is compatible with Linux, Mac and Windows (using Windows Subsystem for Linux) operating systems. This interface collects the IKZF1 index, core Toblerone program, RCH reference values for deletions and functions for extracting relevant reads into a convenient package. This facilitates single sample analysis via a genome aligned BAM file. Once downloaded, it can be easily run as a desktop app. A gene and cohort must be selected (currently only the IKZF1 gene and the RCH B-ALL cohort) along with an indexed genome aligned RNA-seq BAM file. The relevant IKZF1 reads are extracted and mapped with Toblerone to the IKZF1 deletion transcriptome. Finally, the results are displayed in an interactive format along with the high, medium and low confidence thresholds for clinical deletions, as well as outlier information for those without absolute values of reference. Further cohorts can be added to the app in future.

## Data and Code availability

RNA-seq for IKZF1 99 primary RCH cohort samples and selected relapse samples is available from the European Genome-Phenome Archive (accession number EGAS00001004212). Data from the published CD22 resistance patients is publicly available in the Short Read Archive, BioProject ID PRJNA764243. The original template Rust pseudoaligner from 10XGenomics is on GitHub: https://github.com/10XGenomics/rust-pseudoaligner. Source code for the Toblerone adaption is available at: https://github.com/oshlack/toblerone along with instructions for compilation and usage. Source code for the app is available at: https://github.com/Oshlack/TobleroneApp and binaries can be downloaded directly from http://oshlacklab.com/TobleroneApp/

## Acknowledgements

- Tumour samples were made available by the Children’s Cancer Centre Tissue Bank at the Murdoch Children’s Research Institute and The Royal Children’s Hospital (www.mcri.edu.au/childrenscancercentretissuebank), made possible through generous support by CIKA (Cancer In Kids @ RCH), The Royal Children’s Hospital Foundation and the Murdoch Children’s Research Institute.
- This work was supported by NHMRC project grant GNT1140626 to AO and PGE. AO is supported by NHMRC Investigator Grant GNT1196256.

