## Supplemental Figures for "Toblerone: detecting exon deletion events in cancer using RNA-seq"

### Toblerone Supplementary Figures

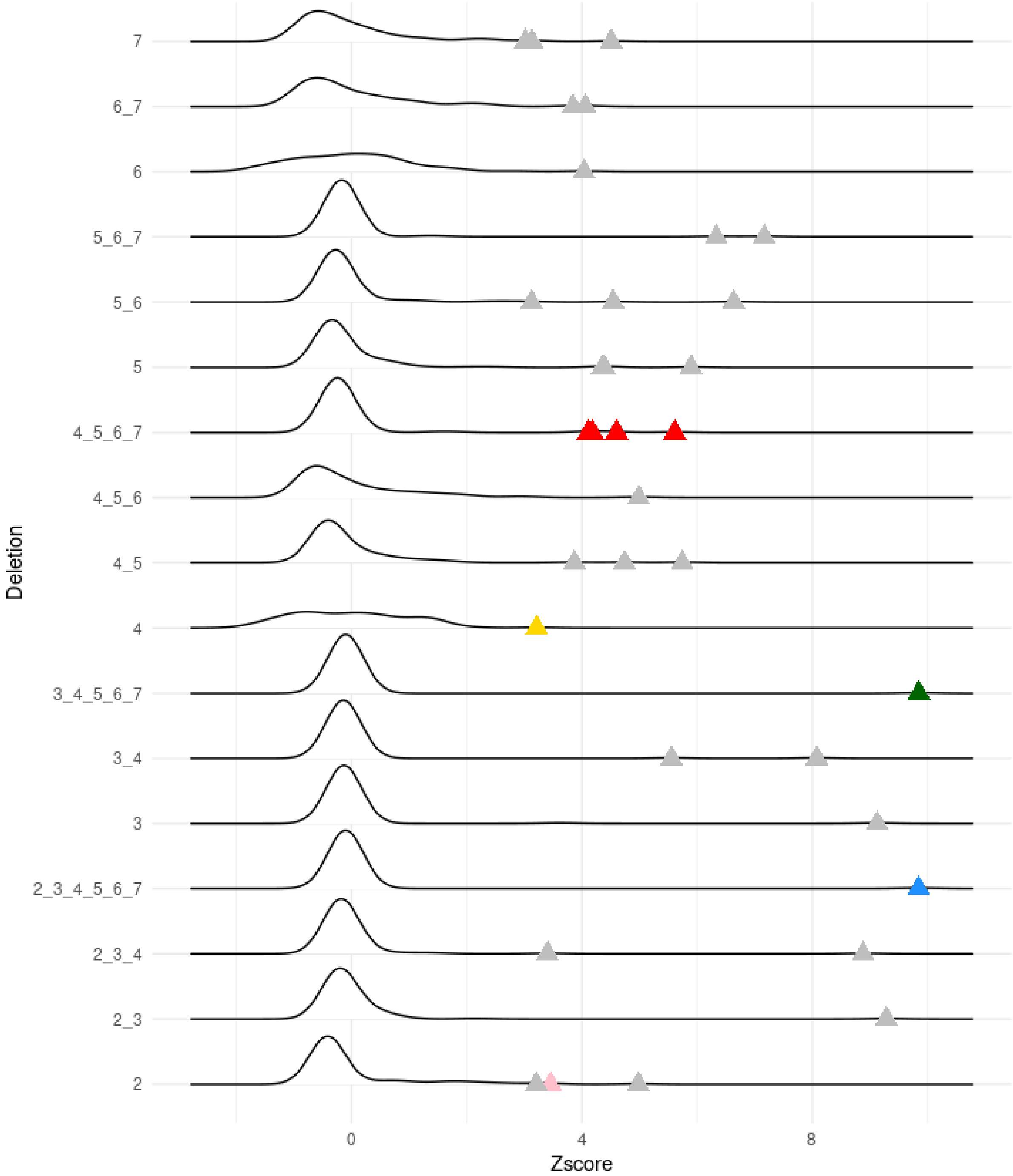


Supplementary Figure 1. Z-score distributions for each IKZF1 deletion with at least one Z-score outlier ≥ 3 with outliers shown as triangles coloured as in Figure 1A. No minimum UC count was applied to define an outlier.

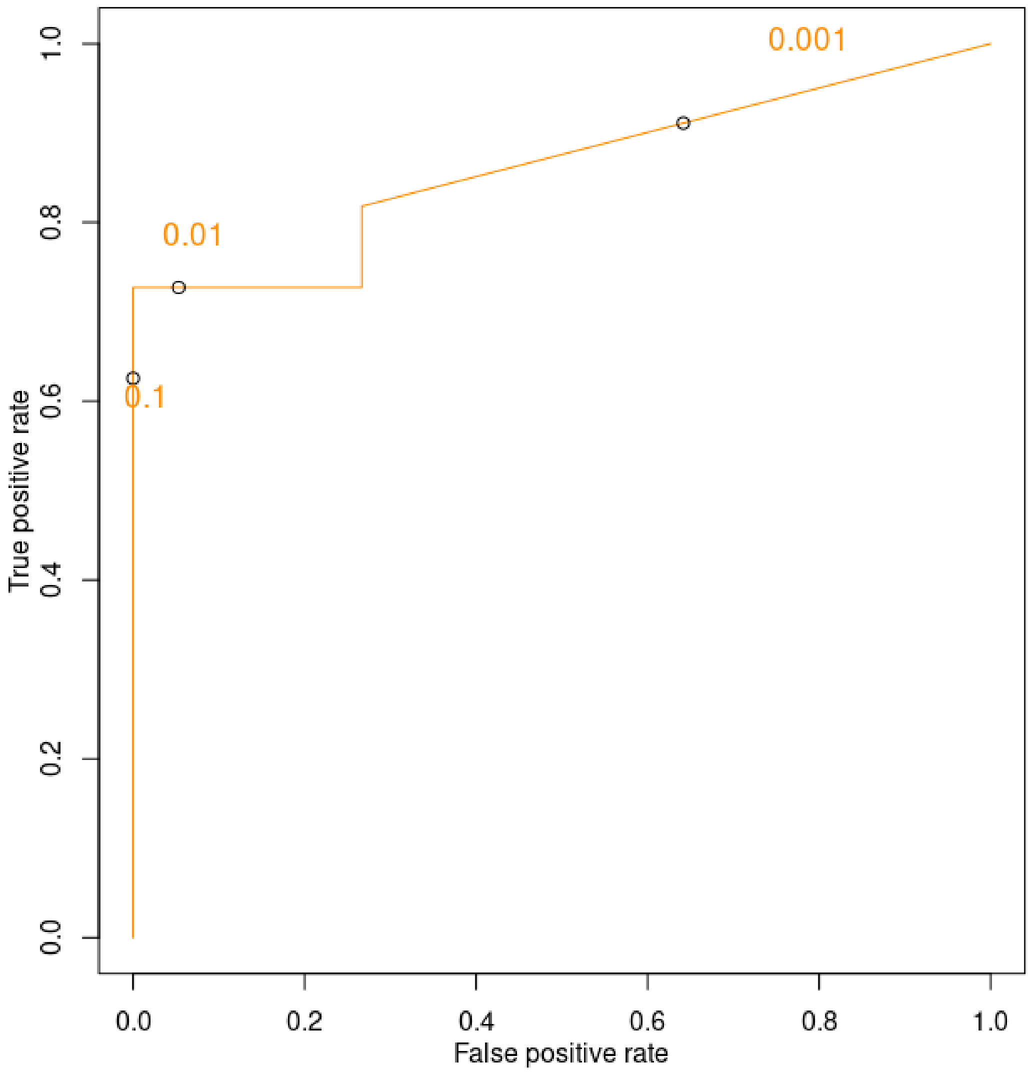


Supplementary Figure 2. A ROC curve of Toblerone Scaled Proportions using the MLPA verified true positives and true negatives for IKZF1 del 4-7 in 26 tested samples (11 True Positive, 15 True Negative ). The high (0.3), medium (0.015) and low (0.001) confidence thresholds are annotated. Area under the ROC is 0.860.

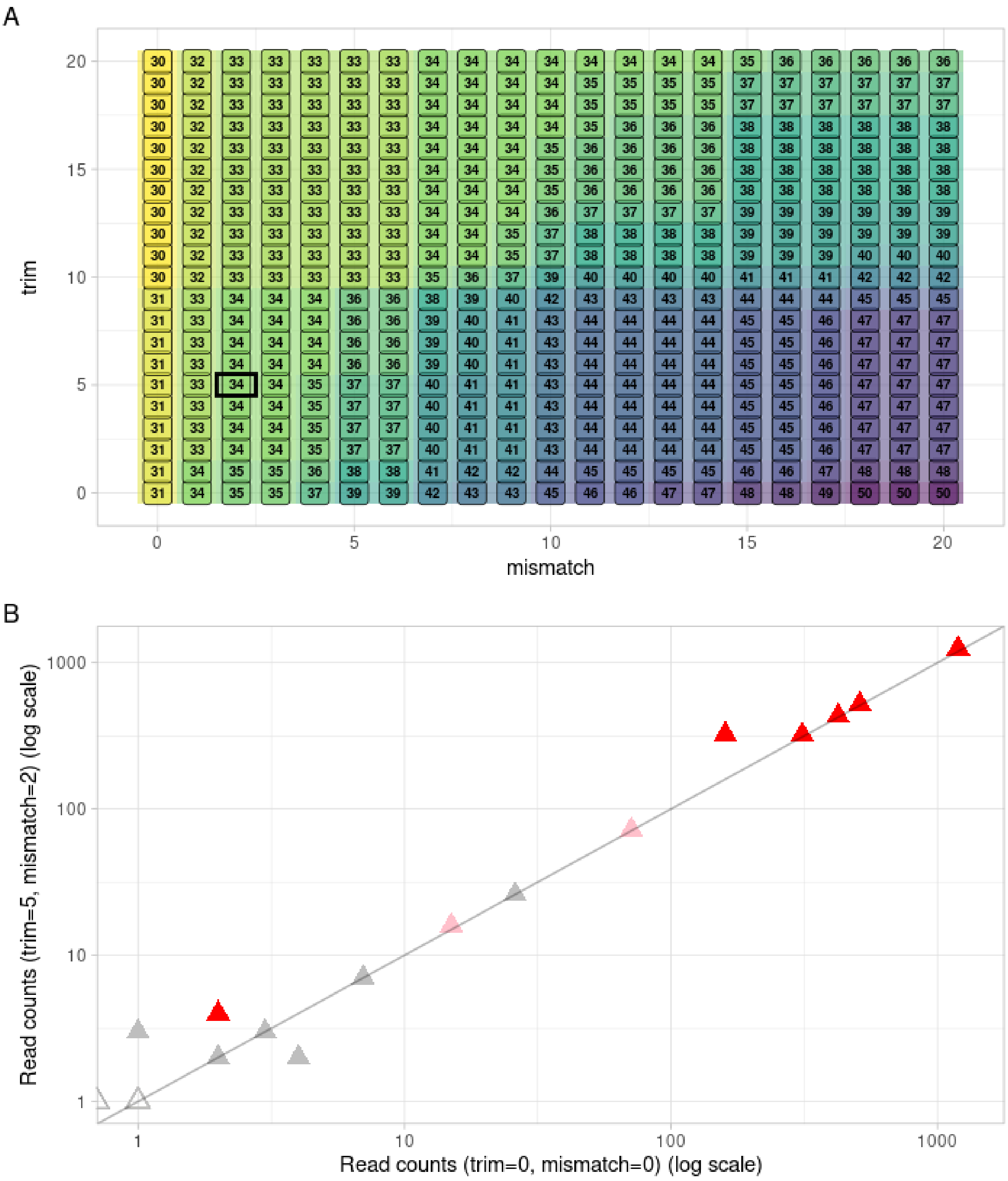


Supplementary Figure 3. Effect of parameter selections to results. A) Number of samples with 1 or more IKZF1 del4_7 counts for varying parameters of trim and mismatch, with default selected parameters for Toblerone boxed. B) Effect of default parameters (trim 5, mismatch 2) on UC counts for IKZF1 del 4_7 deletions, annotated by colour and fill as above, shown on a log scale.

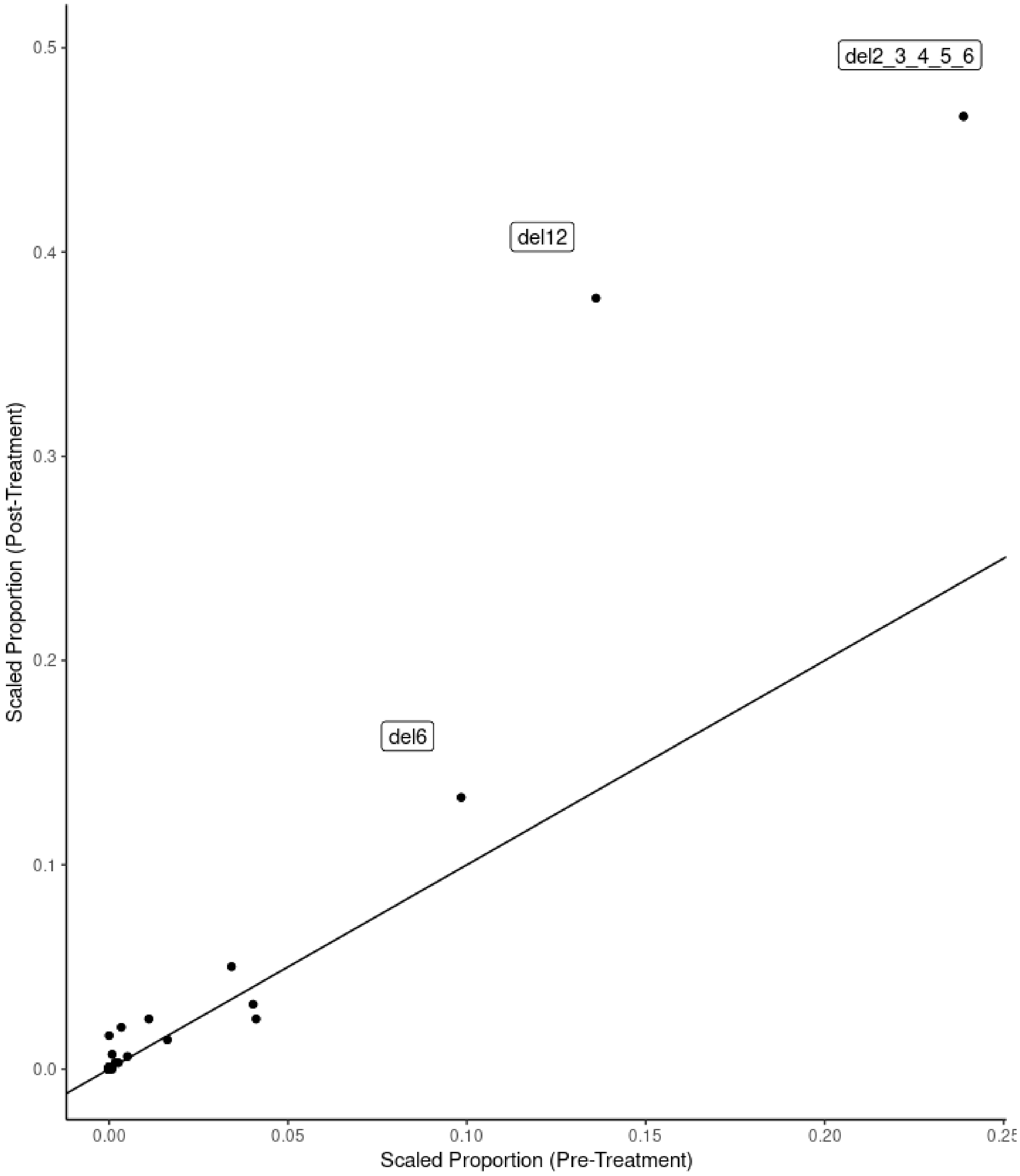


Supplementary Figure 4. Scaled proportions for patient PABDRV in [Zheng et al.](#) before (x-axis) and after (y-axis) treatment with inotuzumab. Line showing equal proportions in black. Notable differences post CAR T-cell therapy include deletion of exon 6, exon 12, and exon 2 through 6.
